# Animal model and therapeutic prospects for left ventricular hypertrophy and associated renal complication

**DOI:** 10.1101/638353

**Authors:** Ashfaq Ahmad, Zainab Riaz, Munavvar Abdul Sattar, Safia Akhtar Khan, Edward James John, Sumbal Arshad, Syed Tahir Abbas Shah, Muhammad Arshad Rafiq, Maleeha Azam, Raheel Qamar

## Abstract

Cardiac and renal dysfunction often co-occur and considerably worsen the prognosis leading to difficulty in therapy in left ventricular hypertrophy (LVH). The aim of this study was to elucidate changes in expression of human ortholog genes of hypertension, vascular and cardiac remodeling and hypertensive nephropathy phenotypes under normal, disease and gasotransmitter, H_2_S (hydrogen sulphide) and NO (nitric oxide) and combined (H_2_S+NO), treatment of rat myocardium and renal tissues. LVH rat models were generated and were treated with H_2_S and NO. Relative gene expression was quantified. Heart and renal physical indices were significantly modified under individual as well as combined H_2_S+NO treatment in control and LVH rats. Expression analysis revealed, hypertension, vascular remodeling genes *ACE, TNFα* and *IGF1*, mRNAs to be significantly increased (P<0.05) in myocardia and kidneys of LVH rats, while individual and combined H_2_S+NO treatment reduced gene expression to normal/near to normal values. The cardiac remodeling genes *MYH7, TGFβ, SMAD4* and *BRG1* expression were significantly up-regulated (P<0.05) in the myocardia of LVH and combined H_2_S+NO treatment, which recovered the normal/near to normal expression more effectively as compared to individual treatments. Interestingly, individual as well as combined H_2_S and NO treatment significantly decreased *PKD1* expression in renal tissue, which was significantly up-regulated in LVH rats (P<0.05). The reduction in hemodynamic parameters and cardiac indices as well as alteration in gene expression on treatment in LVH rat model indicates important therapeutic potential of combined treatment with H_2_S+NO gasotransmitters in hypertension and cardiac hypertrophy associated with renal complications.

## Introduction

Left ventricular hypertrophy (LVH) develops as an adaptive response to increased hypertension induced cardiac workload and volumetric overload. The hypertrophied myocardium adversely affects cardiac function and is a major factor contributing to heart failure [1]. Cardiac and renal dysfunction often co-occur and considerably worsen the prognosis; renal dysfunction can be causative and/or a complication of cardiovascular disease [1]. Physiological cardio-renal interactions can be explained in terms of systemic hemodynamics and renal-body fluid system for blood pressure control [2]. Activation of the sympathetic nervous system and renin-angiotensin system (RAS) in response to decreased cardiac output causes volume expansion and restores renal blood flow [3]. Renal control of blood pressure has a near infinite feedback gain principle such that renal failure, as a consequence of heart failure, causes sodium and water retention, which further exacerbates the cardiac failure [2]. Hypertension induced LVH is associated with vascular remodeling, mediated by vasoactive molecules and endothelial dysfunction as prominent pathological features [4]. Moreover, gene expression changes in the myocardium following stimuli such as pressure overload results in expression of alternate isoform of sarcomeric proteins, which is a major factor contributing to cardiac remodeling and hypertrophied growth of the adult heart muscle [5].

The gasotransmitters hydrogen sulphide (H_2_S) and nitric oxide (NO) are increasingly being studied for their therapeutic potential in treating cardiovascular disease [6,7]. Endogenously H_2_S is produced by pyridoxal-5’-phosphate (PLP)-dependent enzymes [8], while NO is generated by nitric oxide synthases [9]. Several studies have demonstrated the pivotal role of H_2_S in protecting against cardiac hypertrophy by reducing oxidative stress [10], myocardial apoptosis [11], hypertrophy and collagen content [12] and down-regulation of angiotensin II levels [13]. The development of cardiac hypertrophy in mice lacking nitric oxide synthases [14] and the attenuation of LVH in hypertensive rats following treatment with L-arginine to improve NO production [15] highlights the therapeutic role of NO in ameliorating LVH. Recently, a hypothetical model proposed by Yong et al. [16] and Nagpure et al. [17] have shown the existence of interaction between H_2_S and NO, due to the production of intermediate HSNO (thionitrous acid) and HNO (nitroxyl) gases [16], the synergistic effect elicited by H_2_S, NO and these intermediate product is proposed to lead to cardio protection, maintenance of vascular tone and reduced oxidative stress. Recent studies have also shown that the pyridoxal-5’-phosphate (PLP)-dependent enzymes that produce H_2_S are expressed in the kidney [18], where H_2_S shows a protective role against chronic renal failure [19], renal ischemia/reperfusion injury [20] and control renal tubular and vascular function [21]. NO plays a significant role in regulating renal hemodynamics and tubular function [22] and, therefore, maintenance of arterial pressure and vascular volume [23]. This notion is further supported by the implication of NO deficiency in the progression of hypertension and renal diseases [24]. However, the effects of modulatory control of combined therapy with gasotransmitters H_2_S and NO, for the treatment of cardiovascular and renal disease remains largely unexplored.

In the present study we investigated the modulation of the expression of specific genes in response to individual and combined treatment of H_2_S and NO gasotransmitters in the myocardium and renal tissue of control and LVH rat models. The studied genes are known to be involved in hypertension, vascular and cardiac remodeling and hypertensive nephropathy in human.

## Materials and Methods

### Study approval

Animal experimentation protocols used in this study were approved by the Animal Research and Service Centre “ARASC” under the Animal Ethics Committee, Universiti Sains Malaysia “AECUSM” (approval number: 2012/76/364) and Ethic Review Board of the Department of Biosciences, COMSATS Institute of Information Technology, Islamabad, Pakistan.

### Experimental Animals

Male Wistar-Kyoto (WKY) rats with body weight in the range of 180-200g were procured from ARASC and acclimatized for 5 days in the new environment with food and water available ad libitum. Isoprenaline (5mg/kg S/C every 72 hrs) was administered subcutaneously as 5 injections and caffeine (62mg/L) was given in drinking water to the WKY rats for 14 days to induce LVH [25]. The H_2_S donor NaHS (56μM/kg) was administered exogenously in daily intraperitoneal injections for 5 weeks [26], beginning 3 weeks prior to isoprenaline and caffeine administration, while the control groups received saline injections in a similar manner. L-arginine (1.25g/L) was given in drinking water as NO donor for 5 weeks [27].

Seventy two rats were divided into two groups, one for the cardiovascular investigations and renal functional study in acute experiment and the other group for molecular expression studies of the myocardium and renal tissues. Both groups were further divided into 8 sub-groups: Control, LVH, Control-H_2_S (NaHS treated control), Control-NO (L-arginine treated control), Control-H_2_S+NO (NaHS and L-arginine treated control), LVH-H_2_S (NaHS treated LVH), LVH-NO (L-arginine treated LVH) and LVH-H_2_S+NO (NaHS and L-arginine treated LVH); both H_2_S and NO were given separately to the same animal groups (n=48 i.e. 6 rats per group) for the acute studies and n=24 i.e. 3 rats per group for the molecular expression studies.

### H_2_S and NO plasma concentration measurement

At the end of the treatment period, tail vein blood sample was taken from each rat and subjected to centrifugation at 2000 x g for 10 minutes [25]. Plasma H_2_S concentration was measured using a standard procedure [21,26]. NO level in the plasma was measured using a laboratory kit (NJJC Bio Inc., Nanjing, China) by following the manufacturer’s instructions.

### Acute experiment

The acute experiment (on anesthetized non-behaving animals) was performed according to previously reported protocol [25]. The rats were made to fast overnight, by removing food from their cages the night before the experiment, and anaesthetized by intraperitoneally administering 60mg/kg pentobarbitone sodium (Nembutal; CEVA Sante Animale, Libourne, France). A cannula was introduced into the trachea to facilitate ventilation. Following this the right carotid artery was cannulated (PP50, Portex, Kent, UK) and the cannula was connected to a pressure transducer (model P23 ID Gould, Statham Instruments, UK), which was in turn connected to a data acquisition system (PowerLab®, ADInstrument, Australia) to continuously monitor mean arterial pressure (MAP). The left jugular vein was cannulated (PP50, Portex, Kent, UK) to allow infusion of maintenance doses of anesthesia and saline as required. A mid-line abdominal incision exposed the left kidney, which was covered with saline soaked pads to prevent drying. A cannula (PP50, Portex, Kent, UK) was inserted into the iliac artery and attached to another fluid filled pressure transducer connected to the PowerLab system to measure iliac mean blood pressure. A laser Doppler flow probe (AD Instruments, Australia), positioned on the outermost layer of the left kidney cortex was used to record the renal cortical blood perfusion (RCBP). The MAP and RCBP were measured continuously for an hour before the rats were euthanized with an overdose of anesthetic and the hearts were harvested post-mortem, dried and weighed for heart and left ventricle (LV) indices to observe the induction of LVH and how treatment effects the regression of LVH. Internal diameter of the LV chamber and myocardium thickness were measured using vernier caliper according to previously reported protocol [28]. Pulse wave velocity (PWV) was measured by taking the propagation time of the pulse wave from Power Lab data and propagation distance was measured manually by putting a thread from the insertion point of the carotid artery cannula to that of the iliac artery cannula [29].

### RNA extraction and expression analysis in myocardia and renal tissues of control, LVH, H_2_S and NO treated rats

Target tissues were extracted, disrupted and homogenized as previously described [30]. Briefly, following cervical dislocation, heart muscle and renal cortex were harvested and preserved in RNAlater solution (Ambion, Life Technologies Corporation, Carlsbad, CA) at 4°C. RNAZap (Ambion, Life Technologies Corporation, Carlsbad, CA) solution was used to wash all equipment in order to maintain RNA integrity and prevent contamination. Total RNA was extracted using TRIzole reagent (Ambion, Life technologies, Carlsbad, CA) following the manufacturer’s guidelines; after consecutive homogenization, washing and elution, total RNA was extracted and quantified for yield and purity using a microplate reader (Bio Tek Instrument. Inc., Winooski, VT). Total RNA was converted to cDNA using High-Capacity RNA-to-cDNA Kit (Applied Biosystems, Foster City, CA) according to the manufacturer’s protocol using a StepOne Plus real-time polymerase chain reaction (RT-PCR) (Applied Biosystems, Foster City, CA). Relative gene expression of hypertension and vascular remodeling genes; angiotensin I converting enzyme (*ACE*; NM_012544.1), prostacyclin synthase (*PTGIS*; NM_031557.2), regulator of G protein signaling 5 (*RGS5*; NM_019341.1), tumor necrosis factor α (*TNFα*; NM_012675.3), insulin-like growth factor 1 (*IGF1*; NM_001082478.1), were analyzed in heart and renal tissues, while cardiac remodeling genes; transforming growth factor β1 (*TGFβ1*; NM_021578.2), myosin heavy chain 7 (*MYH7*; NM_017240.2), brahma-related gene 1 (*BRG1*; NM_134368.1), SMAD family member 4 (*SMAD4*; NM_019275.3), were studied in the cardiac tissues (Table 1). To determine the effect of H_2_S, NO and H_2_S+NO treatment on renal complication in LVH rats, expression of polycystic kidney disease gene; polycystic kidney disease 1 (*PKD1*; NM_001106352.1) was determined in renal tissue (Table 1), while glyceraldehyde 3-phosphate dehydrogenase (*GAPDH*; NM_017008.4) was used as the housekeeping gene internal control. Primers for RT-PCR were designed using an online primer-designing tool (https://eu.idtdna.com/site at Integrated DNA Technologies, San Diego, CA). RT-PCR reactions were performed on RNA extracted from 3 animals each from the 8 groups (Control, LVH, Control-H_2_S, Control-NO, Control-H_2_S+NO, LVH-H_2_S, LVH-NO and LVH-H_2_S+NO) and each heart and kidney sample was further analyzed in triplicate using maxima SYBR Green/ROX qPCR Kit (Thermo Scientific, Waltham, MA) according to the manufacturer’s instructions on the AB1 StepOne RT-PCR system (Ambion, Foster City, CA). Relative quantification of target gene along with internal control was calculated using the comparative cycle threshold method (2^-ΔΔCT^).

**Table 1.**
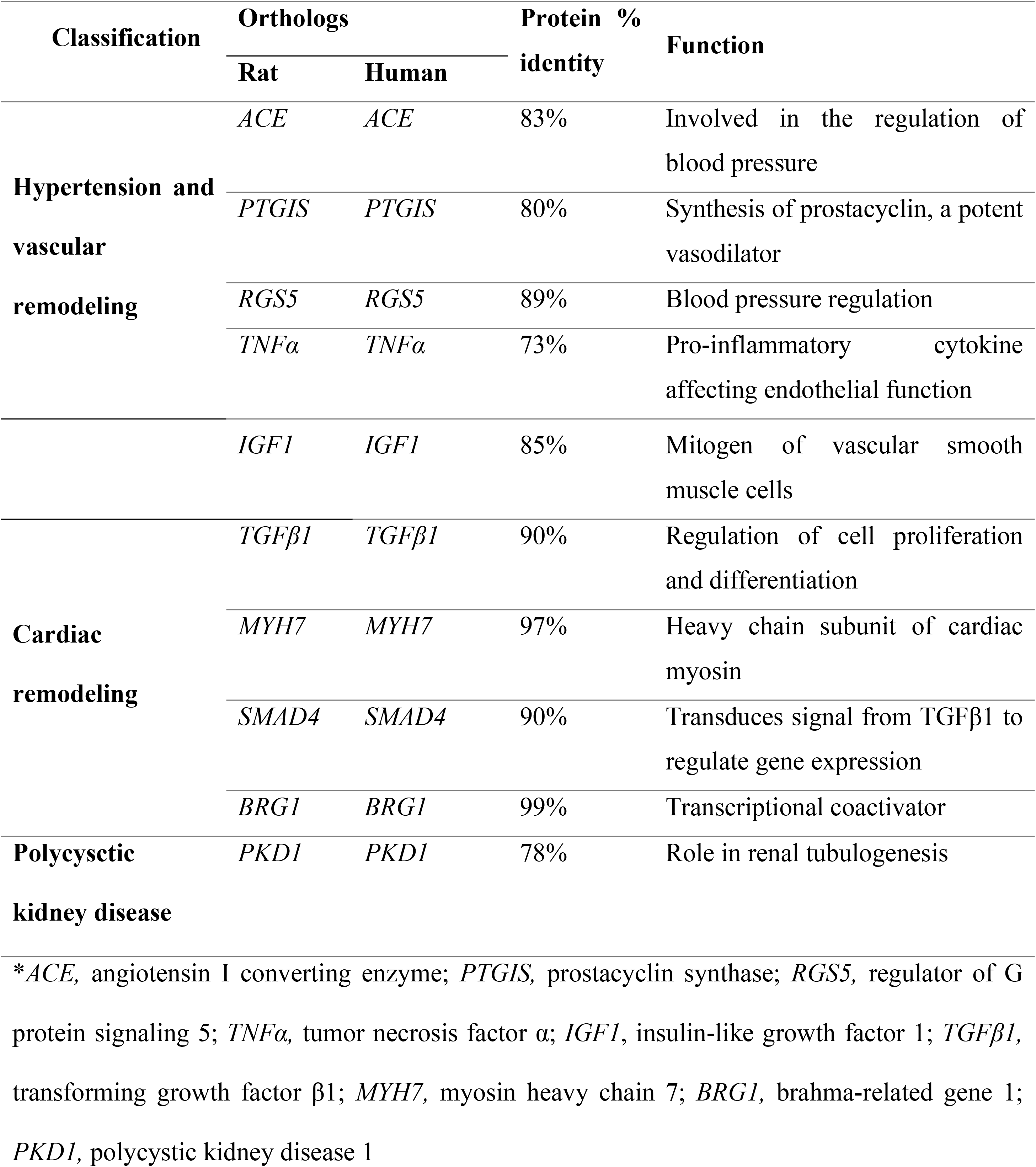
Orthologous genes, % identity of protein (Ensembl 2016) and cellular function.

### Statistical Analysis

Statistical analysis on the systemic hemodynamic data, LV and heart indices and gene expression data was performed using a one way analysis of variance (ANOVA), proceeded by Tukey’s honestly significant difference (HSD) post hoc test using either Origin 2017 (OriginLab, Northampton, MA) or Graph Pad Prism (Graph Pad Software, San Diego, CA). Pearson correlation analysis on gene expression data was also performed using Origin 2017 (OriginLab, Northampton, MA). All data are presented as mean ± SEM with significance at P<0.05.

## Results

### Plasma concentration of H_2_S and NO

The concentration of H_2_S and NO was significantly lower in the plasma of the LVH rats compared to control group (P<0.05). H_2_S plasma concentration was significantly increased (P<0.05) on NaHS treatment in control and LVH rats. L-arginine treatment significantly increased (P<0.05) plasma H_2_S concentration in control but not in LVH rats, while combined Larginine and NaHS treatment did not significantly increase H_2_S concentration in LVH, but significantly reduced it in control rats (S1 Fig). NO plasma concentration was significantly increased in individual as well as combined NaHS and L-arginine treatments in both control and LVH rats (P<0.05) (S2 Fig).

### Treatment (H_2_S, NO and H_2_S+NO) ameliorates LVH progression

It was observed that the induction of LVH significantly elevated MAP and PWV as compared to control (P<0.05). Consequently, treatment with H_2_S, NO and H_2_S+NO, significantly reduced MAP by 14%, 20% and 29% and PWV by 25%, 37% and 12%, respectively, in LVH rats (P<0.05). Heart rate (HR) was significantly reduced in LVH rats as compared to control group and H_2_S, NO and H_2_S+NO treatment further reduced HR in LVH rats (P<0.05). RCBP was reduced by ~53bpu in LVH rats as compared to control (P<0.05) and H_2_S, NO and H_2_S+NO treatment resulted in 36%, 42% and 79% higher RCBP compared to untreated LVH rats (P<0.05) (Table 2).

**Table 2.** Systemic hemodynamic parameters of the studied animal groups.

| **Animal Groups** | **Parameters** |  |  |  |
| --- | --- | --- | --- | --- |
|  | MAP (mmHg) | PWV (m/s) | HR (bpm) | RCBP (bpu) |
| Control | 119 ± 1 | 6 ± 0.4 | 314 ± 3 | 150 ± 12 |
| Control-H<sub>2</sub>S | 122 ± 6 | 6 ± 0.2 | 293 ± 3 * | 138 ± 4 |
| Control-NO | 105 ± 1 | 6 ± 0.3 * ψ | 304 ± 14 | 132 ± 3 * |
| **Control-H<sub>2</sub>S+NO** | 117 ± 4 | 7 ± 0.1 * ψ φ | 280 ± 13 * φ | 141 ± 2 |
| LVH | 142 ± 5 * | 8 ± 0.4 * ψ φ σ | 273 ± 2 * ψ φ | 97 ± 6 * ψ φ σ |
| LVH-H<sub>2</sub>S | 122 ± 3 # | 6 ± 0.3* # | 247 ± 10 * # | 132 ± 5 * # |
| LVH-NO | 114 ± 3 # | 5 ± 0.3* # | 257 ± 7 * | 138 ± 9 # |
| LVH-H<sub>2</sub>S+NO | 101 ± 5 * # λ | 7 ± 0.2 # λ π | 228 ± 5 * # λ π | 174 ± 14 * # λ π |

\*Data are expressed as mean ± SEM and significance taken at P<0.05. \* significant vs. control group; ψ significant vs. control-H<sub>2</sub>S; φ significant vs. control-NO; σ significant vs. control-H<sub>2</sub>S+NO; # significant vs. LVH; λ significant vs. LVH-H<sub>2</sub>S; π significant vs. LVH-NO.
†MAP, mean arterial pressure; PWV, pulse wave velocity; HR, heart rate; RCBP, renal cortical blood perfusion; Control-H<sub>2</sub>S, NaHS treated control group; Control-NO, L-arginine treated control group; Control-H<sub>2</sub>S+NO, NaHS and L-arginine combined treated control group; LVH, left ventricular hypertrophy induced rats; LVH-H<sub>2</sub>S, NaHS treated LVH rats; LVH-NO, L-arginine treated LVH rats; LVH-H<sub>2</sub>S+NO, NaHS and L-arginine combined treated LVH rats.

As a measure of heart and kidney physical parameters, heart, LV index and renal indices were observed to increase by about 46%, 53% and 17% (P<0.05), respectively, in LVH rats as compared to control group and treatment with H_2_S and NO reduced these parameters; combined treatment reduced heart, LV index and renal indices by 41%, 22% and 9% (P<0.05), respectively. Moreover, induction of LVH increased the myocardium thickness by 1.6mm and reduced LV chamber diameter by ~2mm and H_2_S and NO treatment decreased myocardium thickness and increased LV chamber diameter (P<0.05). H_2_S+NO treatment reduced thickness of the myocardium by 36% and increased LV chamber diameter by 43% in LVH rats (Table 3).

**Table 3.** Heart and kidney physical parameters of the studied animal groups.

| Animal Groups | Heart and kidney physical indices |  |  |  |  |
| --- | --- | --- | --- | --- | --- |
|  | Heart index (%) | LV index (%) | Kidney index (%) | Myocardium thickness (mm) | LV chamber diameter (mm) |
| Control | 0.26 ± 0.003 | 0.15 ± 0.004 | 0.29 ± 0.003 | 1.64 ± 0.07 | 5.09 ± 0.01 |
| **Control-H<sub>2</sub>S** | 0.27 ± 0.01 | 0.19 ± 0.006 * | 0.32 ± 0.01 * | 1.80 ± 0.05 * | 4.2 ± 0.16 * |
| Control-NO | 0.24 ± 0.004 | 0.17 ± 0.002 | 0.30 ± 0.01 | 1.81 ± 0.07 * | 5.85 ± 0.10 * ψ |
| **Control-H<sub>2</sub>S+NO** | 0.30 ± 0.002 * φ | 0.20 ± 0.004 * φ | 0.30 ± 0.01 | 2.05 ± 0.08 * ψ φ | 4.42 ± 0.18 * ψ φ |
| LVH | 0.38 ± 0.007 * ψ φ σ | 0.23 ± 0.002 * ψ φ σ | 0.34 ± 0.007 * | 3.29 ± 0.04 * ψ φ σ | 2.85 ± 0.04 * ψ φ σ |
| **LVH-H<sub>2</sub>S** | 0.34 ± 0.01 * # π | 0.21 ± 0.001 * | 0.32 ± 0.01 * | 2.36 ± 0.05 * # | 5.38 ± 0.05 * # |
| LVH-NO | 0.30 ± 0.002 * # λ | 0.21 ± 0.003 * | 0.31 ± 0.007 # | 2.28 ± 0.09 * # λ | 4.80 ± 0.11 * # λ |
| **LVH-H<sub>2</sub>S+NO** | 0.27 ± 0.004 # λ π | 0.18 ± 0.003 * # λ π | 0.31 ± 0.007 # | 2.09 ± 0.04 * # λ π | 4.09 ± 0.07 * # λ π |

\* Data are expressed as mean ± SEM and significance taken at P<0.05. \* significant vs. control group; ψ significant vs. control-H<sub>2</sub>S; φ significant vs. control-NO; σ significant vs. control-H<sub>2</sub>S+NO; # significant vs. LVH; λ significant vs. LVH-H<sub>2</sub>S; π significant vs. LVH-NO.

### Expression modulation of hypertension and vascular remodeling genes under H_2_S, NO, and H_2_S+NO treatment

There was a ~2 fold increase in *ACE* expression in the myocardium and ~7.5 fold increase in *ACE* expression in the renal tissue of LVH rats as compared to control (P<0.05). In both, heart and kidney, H_2_S treatment in control significantly up-regulated *ACE* expression while H_2_S treatment in LVH rats significantly reduced *ACE* expression (both P<0.05). NO treatment significantly down-regulated *ACE* expression in both control and LVH group myocardia and kidneys (both P<0.05). Combined treatment with H_2_S+NO significantly down-regulated (P<0.05) *ACE* in kidney of LVH rats, however, down-regulation of *ACE* expression was not significant in heart (P>0.05) (Fig 1A). *PTGIS* expression increased by 1.5 fold in heart tissue of LVH rats as compared to control and H_2_S and H_2_S+NO treatment showed significant down-regulation, while NO treatment increased expression in heart tissue of LVH rats (P<0.05). Contrarily, in renal tissue *PTGIS* expression was significantly down-regulated in LVH group as compared to control, however, individual as well as combined H_2_S and NO treatment up-regulated *PTGIS* in LVH rats (P<0.05) (Fig 1B). *RGS5* was 0.2-0.3 fold down-regulated in both heart and renal tissues of LVH as compared to control group (P<0.05) but there was no notable significant change in gene expression of *RGS5* after H_2_S, NO and H_2_S+NO treatment in LVH rats (Fig 1C).

**Fig 1.**
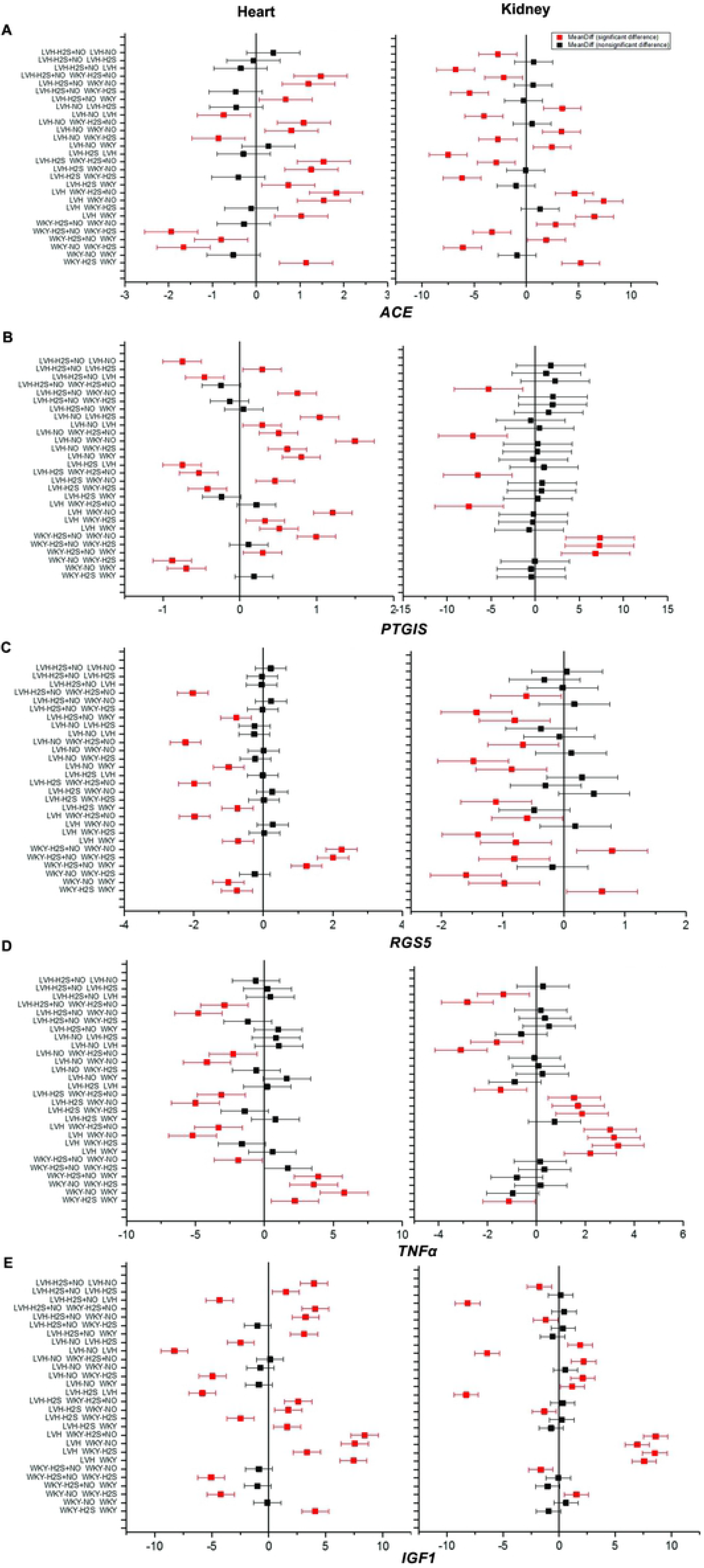
Graphical representation of relative gene expression of hypertension, vascular remodeling genes. Graph showing the relative gene expression in control, left ventricular hypertrophy (LVH) and treated groups in cardiac and renal tissues of hypertension, vascular remodeling genes. Relative gene expression of (A) *ACE*, (B) *PTGIS*, (C) *RGS5*, (D) *TNFα*, (E) *IGF1*, in cardiac and renal tissues of control, LVH, H_2_S and NO treated and H_2_S+NO combined treated rats.

*TNFα* was significantly up-regulated in both renal and heart tissues of LVH WKY as compared to control (P<0.05), while H_2_S, NO and H_2_S+NO treatment significantly down-regulated *TNFα* expression in kidney (P<0.05), however, it’s expression did not alter in response to H_2_S and H_2_S+NO treatment while NO treatment significantly increased expression in heart tissue of LVH rats (Fig 1D). *IGF1* expression was significantly up-regulated in myocardia and renal tissues of LVH group as compared to control while H_2_S, NO and combined H_2_S+NO treatment significantly down-regulated it normal or near to normal (P<0.05) (Fig 1E).

### Change in cardiac remodeling gene expression in myocardia in response to H_2_S, NO, and H_2_S+NO treatment

Expression of cardiac remodeling genes was studied in the myocardia of LVH rats in response to treatment, *MYH7, TGFβ1, BRG1* and *SMAD4* expression was observed to be significantly up-regulated in myocardia of LVH as compared to control group (all P<0.05) (Fig 2). H_2_S treatment significantly increased *MYH7* expression but there was no significant change in *TGFβ1, BRG1* and *SMAD4* expression in response to H_2_S; however, NO treatment significantly reduced *MYH7* and *BRG1* expression but had no significant effect on *TGFβ1* and *SMAD4* expression in the myocardia of LVH rats. Combined H_2_S+NO treatment did not significantly change the expression of *MYH7, BRG1* and *SMAD4* in the myocardia of the LVH group. However, *TGFβ1* expression was significantly down-regulated (P<0.05) in LVH myocardia treated with H_2_S+NO (Fig 2).

**Fig 2.**
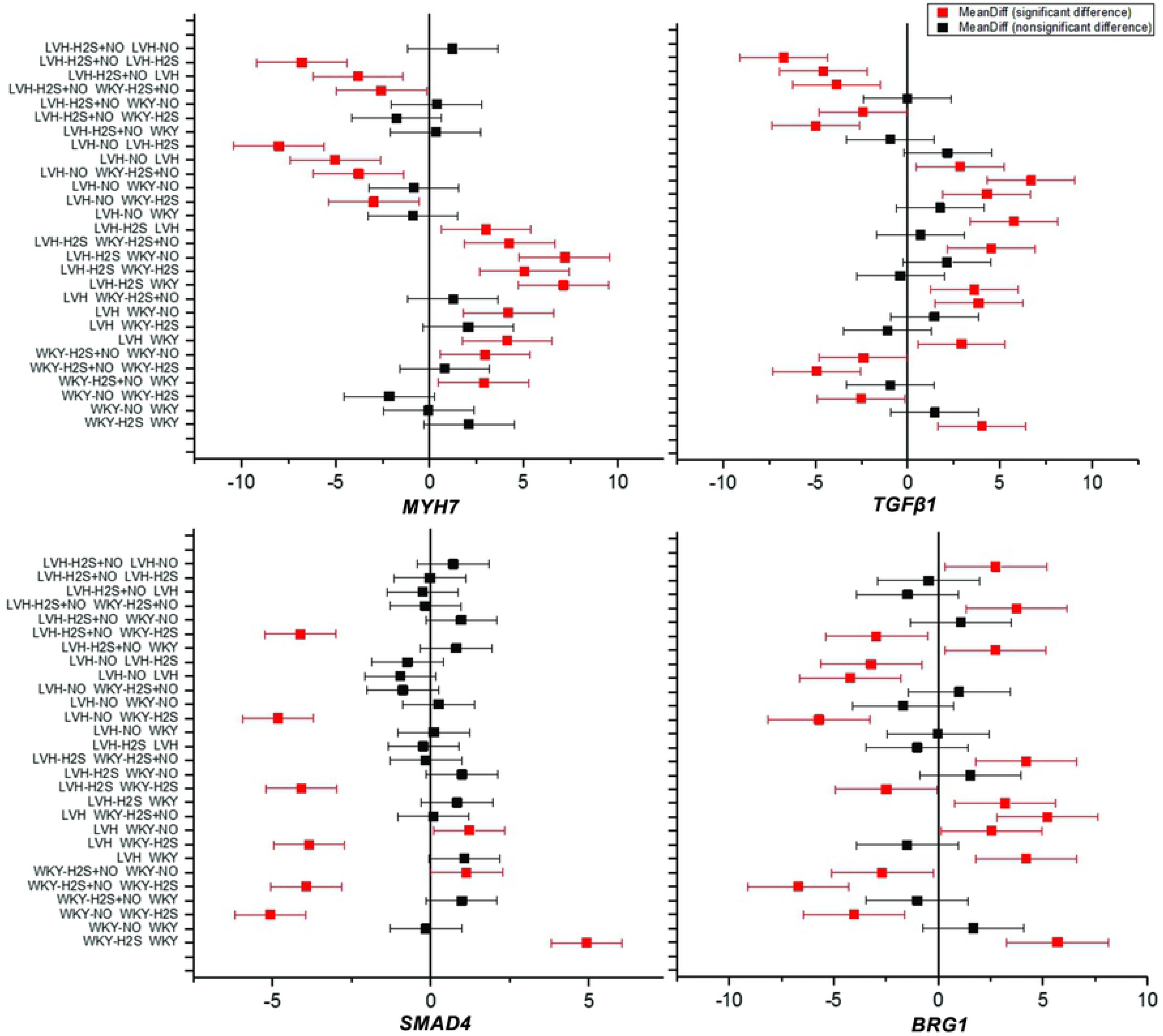
Graph showing the relative gene expression in control, LVH and treated groups of cardiac remodeling genes in cardiac tissue. Relative gene expression of *MYH7, TGFβ1, SMAD4* and *BRG1*, in heart tissues of control, LVH, H_2_S and NO treated and H_2_S+NO combined treated rats.

### Altered *PKD1* expression in renal tissues of H_2_S, NO, and H_2_S+NO treated rats

Hypertension induced LVH is an important complication in polycystic kidney disease (PKD) [31] and to elucidate the effect of H_2_S and NO treatment on hypertensive nephropathy in LVH rats we studied *PKD1* expression in LVH rats before and after treatment. About 3-fold increase in *PKD1* expression was observed in renal tissue of LVH WKY as compared to control group (P<0.05). *PKD1* expression was significantly down-regulated after treatment of LVH rats with H_2_S, NO and H_2_S and NO combined (P<0.05) (Fig 3).

**Fig 3.**
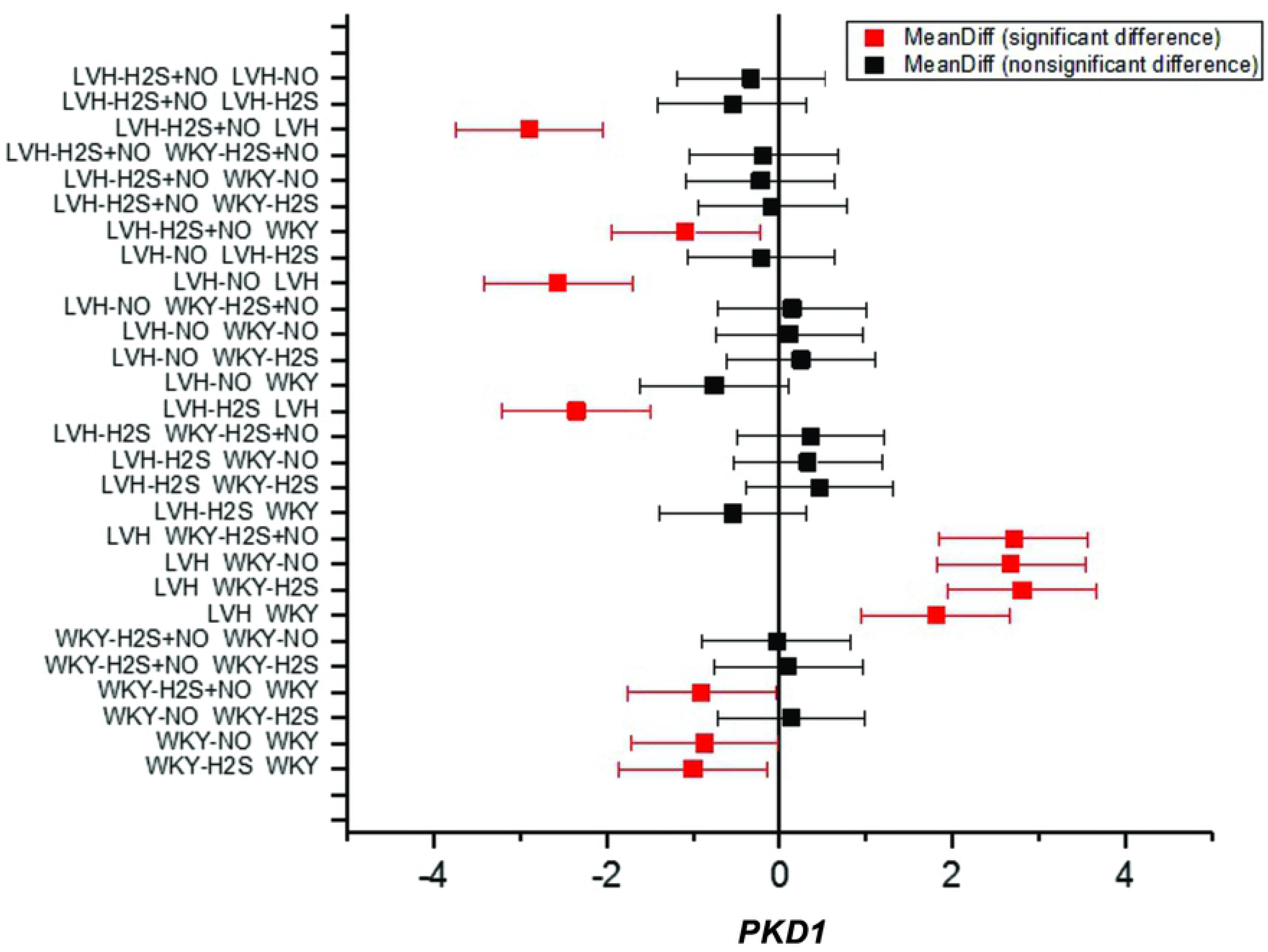
Graph showing the relative gene expression of *PKD1* in control LVH and treated groups in renal tissue.

## Discussion

In this study we aimed to elucidate the modulation in expression of particular orthologous genes upon gasotransmitters, H_2_S and NO, treatment in LVH rats, these genes are known to have a role in hypertension, vascular and cardiac remodeling and hypertensive nephropathy in humans. The significant decrease in MAP, heart index, LV index and myocardium thickness, increase in RCBP, and modulated expression of *ACE, PTGIS, IGF1, TNFα, TGFβ1* and *PKD1* upon exogenous administration of H_2_S+NO, as compared to individual treatments, showed that combined treatment may be more effective in ameliorating LVH and its complications.

Comparison of the pattern of gene expression between all animal groups showed that gene expression was most dissimilar in control and LVH groups, while expression was most similar in control and H_2_S+NO treated LVH groups (Fig 4; S1 Table) thus indicating that H_2_S+NO treatment has promising therapeutic potential in LVH and associated complications.

**Fig 4.**
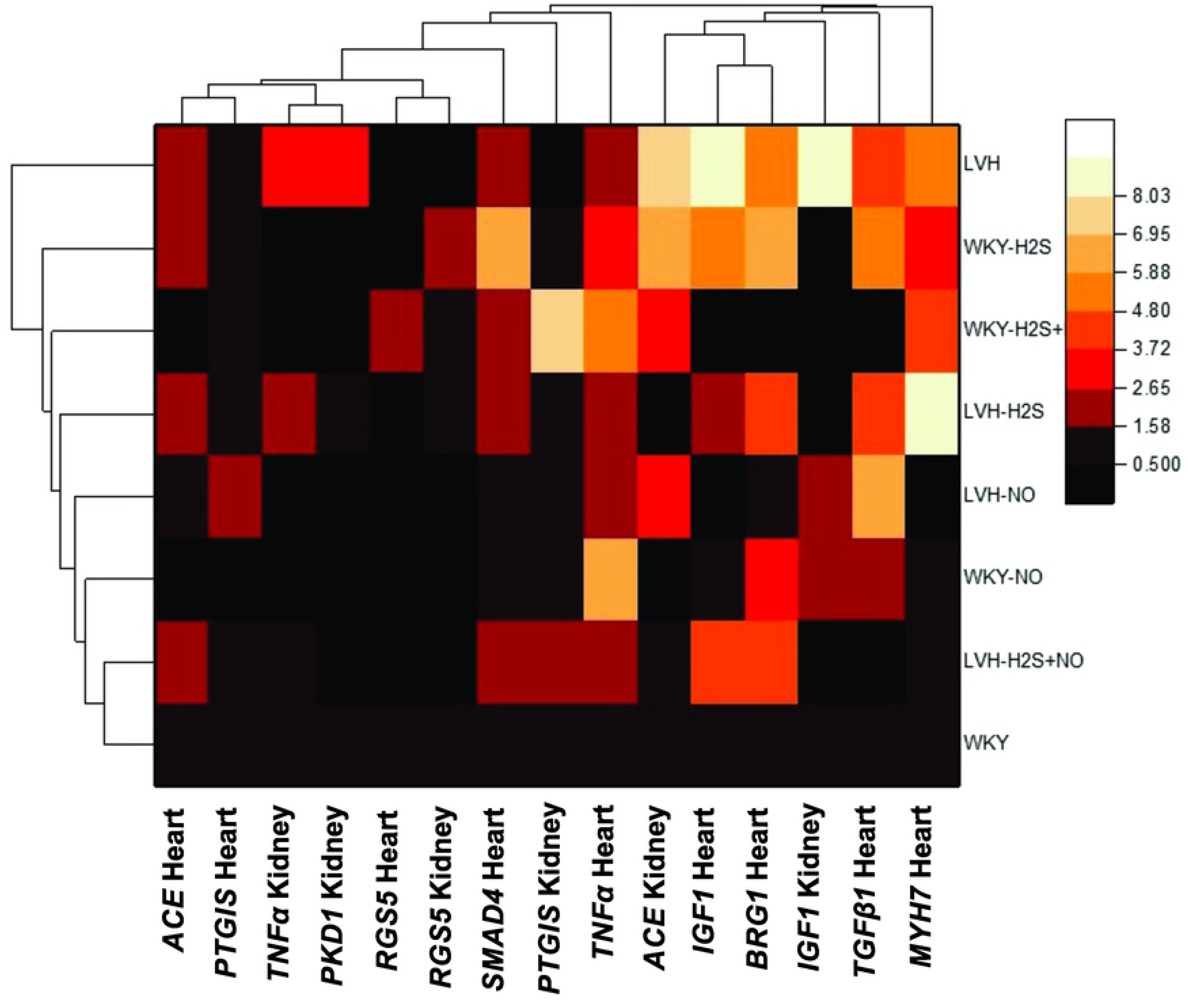
Heatmap showing the comparative relationship of gene expression among studied groups as tested in cardiac and renal tissues.

We previously reported an up-regulation in endothelial nitric oxide synthase (eNOS) and cystathione-gamma lyase (CSE) expression upon exogenous H_2_S administration only [30]. We also reported that individual NO treatment up-regulated eNOS expression in control and LVH rats and CSE expression in control rats, but down-regulated CSE expression in LVH rats [27]. In our previous report on H_2_S+NO treatment we showed that CSE mRNA levels significantly decreased in control but not in LVH rats, while eNOS mRNA levels significantly increased in both LVH and control groups). Extending our study to check H_2_S and NO plasma concentrations, we have found that these changes corresponded to the mRNA levels of their respective synthases. The synthesis of biochemical intermediates and modulation of the respective pathways for NO and H_2_S synthesis by the gasotransmitters may explain the synergistic efficiency of combined H_2_S and NO treatment in ameliorating LVH relative to individual treatments.

The effect of individual and combined treatment with H_2_S and NO was also seen on the expression of genes in LVH WKY rats. Induction of LVH significantly up-regulated *ACE* expression while H_2_S+NO treatment down-regulated *ACE* expression in kidney and heart tissues. Previous studies have reported a two to three fold increase in Angiotensin II (Ang II) levels in isoprenaline induced LVH WKY rats [30,32], which indicates an activation of RAS, resulting in hypertension and arterial stiffness in these LVH models. Being a human ortholog, DNA sequence variations in *ACE* affects circulating ACE concentration and is linked to the development of LVH in humans. The DD genotype of an insertion/deletion (I/D) polymorphism results in elevated ACE plasma levels and is associated with an increased LV mass index [33] and a 2350G>A polymorphism that affects plasma ACE levels is also associated with LVH [34]. Moreover, ACE inhibitors have been shown to be less effective in LVH regression in patients with the DD genotype [35]. The down-regulation of *ACE* expression upon H_2_S+NO treatment in the present study indicates that combined treatment may alleviate hypertension caused by either endogenous or exogenous RAS activation. It was also observed that H_2_S treatment in control WKY rats significantly up-regulates *ACE* expression, which indicates that there may be a difference in the effects elicited by exogenous treatment with gasotransmitters in diseased as opposed to normal settings.

PTGIS catalyzes prostacyclin synthesis from prostaglandin H2 and primarily found in smooth muscle and vascular endothelial cells [36]; prostacyclin suppresses smooth muscle cell proliferation and platelet aggregation and promotes vasodilation [37]. We found a significant up-regulation of *PTGIS* expression in LVH rats as compared to control, while H_2_S+NO treatment down-regulated its expression in LVH rats. This is in line with already reported data in which Champetier et al. [38] observed a 1.9 fold increase in *PTGIS* expression in LV of rats with severe aortic regurgitation, which is associated with left ventricle eccentric hypertrophy, Harding et al. [39] however, reported an anti-hypertrophic nature of prostacyclin and its inhibitory effect on cardiac fibrosis. Genetic changes and epigenetic modifications are known to alter human *PTGIS* ortholog expression where promoter hypermethylation results in reduced expression of *PTGIS* [40]. Based on our findings, there is a significant down-regulation of *PTGIS* in LVH kidney and combined H_2_S+NO treatment up-regulates expression, which is contrary to the gene expression changes in the myocardium. A critical role for prostacyclin in renal development has been shown previously and PTGIS-knockout mice showed severely abnormal glomerular, vascular and interstitial development [41]. There might be a role of epigenetics in differential gene expression between the two tissues, thus indicating the treatment effectiveness also against epigenetic changes in a tissue specific manner.

RGS5 is one of the regulators of G-protein signaling and it is expressed in vascular and cardiac muscle tissues [42] where it serves to inhibit several Gαi- and Gαq-mediated signaling pathways including those that are induced by Ang II [42,43]. *RGS5* human ortholog has been shown to be associated with blood pressure control [44], while, our data has demonstrated a significant reduction in *RGS5* mRNA, hence diminishing its protective role against Ang II mediated signaling, in the heart and kidney tissue of LVH WKY rats as compared to control group and there was no notable change in *RGS5* expression following H_2_S+NO treatment. In addition, *RGS5* expression in heart negatively correlated to the expression of ACE, *TGFβ1* and *IGF1* in the heart in the current study (Fig 4; S1 Table)

IGF1 promotes cell division, it is an antiapoptotic factor in vascular and endothelial cells and promotes cardiac growth and function [45]. Donohue et al. [46] have reported that development of hypertension and hypertrophy was associated with increased *IGF1* mRNA levels in rats. Similarly, Li et al. [47,48] found mRNA and protein levels of the human ortholog to be elevated in idiopathic hypertrophic obstructive cardiomyopathy (HOCM) tissues. Increased IGF1 levels in hypertensive patients as opposed to normotensive individuals is associated with increase in pressure load [49] and is also shown to be positively associated with chronic renal disease [50].

Moreover, promoter polymorphisms in *IGF1* have been shown to be significantly related to LV mass in male athletes [51]. We found *IGF1* expression to be up-regulated in LVH and significant reduction in expression upon H_2_S+NO treatment. TNFα, an inflammatory cytokine has been shown to effect left ventricular remodeling and hypertrophy under hypertension [52] and is a crucial player in the hypertension and cardiac remodeling processes induced by Ang II [53]. TNF-knockout mice have displayed diminished hypertensive response on activation of RAS [54]. The human *TNFα* ortholog by Patel et al. [55] had shown variable expression of *TNFα* in hypertrophic cardiomyopathy (HCM) patients having the AA genotype of *TNFα* −308G>A functional single nucleotide polymorphism, resulting in greater LV mass index development and clinical diagnosis at a younger age. TNFα has been shown to further the progression of cyst formation particularly in an autosomal dominant polycystic kidney disease (ADPKD) genetic background [56]. Our data demonstrates up-regulation of *TNFα* expression in kidney and heart tissues of LVH WKY rats and a significant reduction in TNFα expression in kidney upon NO and H_2_S+NO treatment, moreover, *TNFα* expression showed strong positive correlation with *PKD1* expression (r=0.9) (Fig 4; S1 Table).

The heart muscle responds to cardiac workload and hemodynamic stress by activation of the fetal gene expression program, causing hypertrophic growth [57]. In rats, MYH7 (β-MHC) is the fetal isoform of myosin heavy chain that is shown to be up-regulated in cardiac hypertrophy [58], which is consistent with our findings, where we also observed its higher expression. This *MYH7* overexpression was slightly down-regulated upon LVH regression after H_2_S+NO treatment, although this was not statistically significant. It has been shown by Lowes et al. [59] that β-MHC human ortholog expression increases and expression of α-MHC isoform is reduced in ventricular myocardium in failing hearts. TGFβ1 promotes cardiac remodeling and hypertrophy in response to pressure overload [60]. It has been reported previously that an increase in cardiac *TGFβ1* mRNA in response to AngII infusion in mice and *TNFα* deficient mice failed to elicit this increase [53]. On comparing hypertensive and normotensive individuals, Argano et al. [61] reported a polymorphism in human *TGFβ1* ortholog affecting TGFβ1 sera levels which was found to be associated with hypertension, left ventricular hypertrophy and renal damage. SMAD proteins are transcription activators that are involved in the transduction of TGFβ1 signal [60]. SMAD4 is a crucial part of TGFβ1 signaling pathway [62] and it is reported to induce apoptosis and negative remodeling in hypertrophied myocardia [63]. BRG1 is a chromatin modifying protein that is involved in the switch to fetal MHC expression, promoting hypertrophy [64] and it is reported to be inhibited by H_2_S [65]. In line with this literature, our data demonstrates a significant increase in *TGFβ1, SMAD4* and *BRG1* mRNA in LVH WKY rats. While *TGFβ1* expression was significantly reduced in response to H_2_S+NO treatment, *SMAD4* and *BRG1* expression did not change significantly following treatment. We also observed change in *TGFβ1, MYH7, SMAD4* and *BRG1* expression and moderate to strong correlation between these components (Fig 4; S1 Table).

Another interesting finding of the current study is the modulation in *PKD1* expression in LVH rat, in response to treatment. *PKD1* encodes polycystin-1 which is an integral membrane protein expressed in the renal tubular epithelium [66]. Studies have shown that altered *PKD1* expression may lead to cyst formation in polycystic kidney disease (PKD) [67,68]. Cardiovascular abnormalities such as hypertension and LVH are frequently associated with autosomal-dominant PKD (ADPKD) [69]. We found PKD1 to be ~3-fold up-regulated on induction of LVH in WKY rats and individual treatments as well as combined treatment with H_2_S and NO significantly down-regulated PKD1 expression. Moreover, PKD1 expression showed a strong positive correlation with *ACE, TNFα* and *IGF1* expression in the kidney (Fig 4; S1 Table), the genes that are involved in hypertrophy and vascular remodeling in the heart as well as kidney.

## Conclusion

In summary, in the present study, there was a modulation of expression of the hypertrophy, vascular and cardiac remodeling and hypertensive nephropathy genes in LVH rat model and combined treatment with H_2_S and NO, effectively normalized gene expression in most cases. Human orthologs of these genes have crucial roles in the development of hypertension induced LVH and related complications, which thus highlights a promising therapeutic role for H_2_S and NO combined treatment. Taken together, the present data therefore suggest that combined treatment of H_2_S and NO may be used as a therapeutic intervention for renal and cardiac protection directly and/or indirectly (Fig 5).

**Fig 5.**
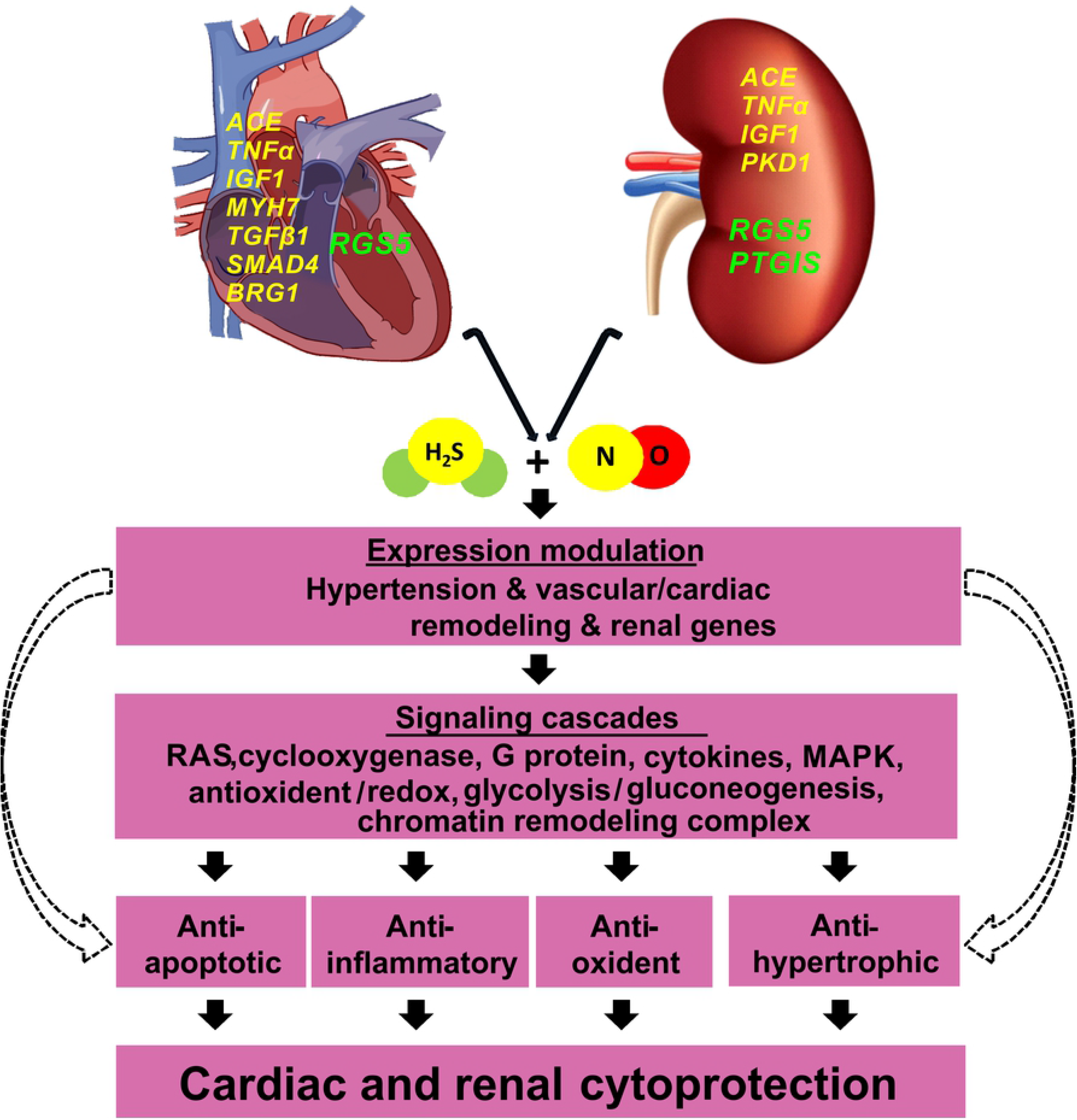
Therapeutic potential of gasotransmitters in left ventricular hypertrophy. Combined treatment of H_2_S+NO has cytoprotective effect in cardiac and renal tissues in left ventricular hypertrophy. The yellow font represents up-regulation while green font represents down-regulation of the studied genes. The dotted arrow is indicative of predicted direct action.

## Acknowledgements

The authors thank all the technician and staff of the labs for facilitation during the course of study.

